# Multi-omics analysis identifies a *CYP9K1* haplotype conferring pyrethroid resistance in the malaria vector *Anopheles funestus* in East Africa

**DOI:** 10.1101/2021.10.21.465247

**Authors:** Jack Hearn, Carlos Djoko Tagne, Sulaiman S. Ibrahim, Billy Tene-Fossog, Leon J. Mugenzi, Helen Irving, Jacob M. Riveron, G.D. Weedall, C.S. Wondji

**Affiliations:** Vector Biology Department, Liverpool School of Tropical Medicine, Pembroke Place, Liverpool L3 5QA, UK; LSTM Research Unit, Centre for Research in Infectious Diseases (CRID), P.O. Box 13591, Yaoundé, Cameroon; Department of Biochemistry, Faculty of Science, University of Bamenda, P.O. Box 39 Bambili, Bamenda, Cameroon; School of Biological and Environmental Sciences, Liverpool John Moores University, Byrom Street, Liverpool L3 3AF, UK

## Abstract

Metabolic resistance to pyrethroids is a menace to the continued effectiveness of malaria vector controls. Its molecular basis is complex and varies geographically across Africa. Here, we used a multi-omics approach, followed-up with functional validation to show that a directionally selected haplotype of a cytochrome P450, *CYP9K1* is a major driver of resistance in *Anopheles funestus*.

A PoolSeq GWAS using mosquitoes alive and dead after permethrin exposure, from Malawi and Cameroon, detected candidate genomic regions, but lacked consistency across replicates. Targeted deep sequencing of candidate resistance genes and genomic loci detected several SNPs associated with known pyrethroid resistance QTLs. The most significant SNP was in the cytochrome P450 *CYP304B1* (Cameroon), *CYP315A1* (Uganda) and the ABC transporter gene ABCG4 (Malawi). However, when comparing field resistant mosquitoes to laboratory susceptible, the pyrethroid resistance locus *rp1* and SNPs around the ABC transporter ABCG4 were consistently significant, except for Uganda where *CYP9K1* P450 was markedly significant. *In vitro* heterologous metabolism assays with recombinant CYP9K1 revealed that it metabolises type II pyrethroid (deltamethrin; 64% depletion) but not type I (permethrin; 0%), while moderately metabolising DDT (17%). *CYP9K1* exhibited a drastic reduction of genetic diversity in Uganda, in contrast to other locations, highlighting an extensive selective sweep. Furthermore, a glycine to alanine (G454A) amino acid mutation located between the meander and cysteine pocket of *CYP9K1* was detected in all Ugandan mosquitoes.

This study sheds further light on the complex evolution of metabolic resistance in a major malaria vector, by adding further resistance genes and variants that can be used to design field applicable markers to better track this resistance Africa-wide.

**Author Summary:** Metabolic resistance to pyrethroids is a menace to the continued effectiveness of malaria vector controls. Its molecular basis is complex and varies geographically across Africa. Here, we used several DNA based approach to associate genomic differences between resistant and susceptible mosquitoes from several field and laboratory populations of the malaria vector *Anopheles funestus*. We followed-up our genomic analyses with functional validation of a candidate resistance gene in East Africa. This gene (*CYP9K1*) is a member of the cytochrome P450 gene-family that helps to metabolise, and thereby detoxify, pyrethroid insecticides. We show that this gene is a major driver of resistance to a specific sub-class of pyrethroid insecticides only, with moderate to no effects on other insecticides used against *Anopheles funestus*. We were able to link resistance in this gene to a mutation that changes the amino acid glycine to alanine that may impact how the protein-product of this gene binds to target insecticides. In addition to demonstrating the biochemical specificity of an evolutionary response, we have broadened the available pool of genes can be used to monitor the spread of insecticide resistance in this species.

## Introduction

Malaria control relies heavily on insecticide-based interventions, notably Long-Lasting Insecticidal Nets (LLINs) incorporating pyrethroid insecticides, and Indoor Residual Spraying (IRS). Together, these interventions are credited with a greater than 70% decrease in malaria burdens since their introduction [1]. However, resistance to insecticides (notably pyrethroids) is threatening the continued effectiveness of these tools. Unless resistance to insecticides is managed, the recent gains in reducing malaria transmission could be lost [2]. Worryingly, several mosquito populations are developing multiple and cross-resistance to a broad range of insecticides, increasing the risks that such populations could be better equipped to rapidly develop resistance to novel classes of insecticides. Therefore, elucidating the genetic basis and evolution of resistance is crucial to design resistance management strategies and prevent malaria resurgence [2].

In the major malaria vector *Anopheles funestus*, metabolic resistance mechanisms are driving resistance to most insecticides, including pyrethroids [3–5]. The molecular basis of this resistance however is diverse and complex across Africa, with different resistance mechanisms spreading, and potentially inter-mixing, from independent origins [6–10]. These mechanisms are driven by extensive genetic variation between regions, preventing the use of existing findings to inform control efforts across the continent. Progress was recently made in this area through the detection of a DNA-marker in the *cis*-regulatory region of the cytochrome P450 *CYP6P9a and CYP6P9b* allowing the design of DNA-based simple PCR assays for detecting and tracking pyrethroid resistance in the field [5, 11]. However, this resistance marker only explains resistance in southern Africa as the genetic basis of pyrethroid resistance, and cross-resistance to other insecticide classes is driven by different genes [5, 9]. This is a major obstacle in designing effective resistance management strategies across the continent, to better control this major malaria vector.

Transcriptomic analyses have successfully been used to detect key genes conferring resistance to insecticides in the principal malaria vectors [4, 5, 12]. Despite large scale whole genome sequencing, it has proven difficult to conclusively associate variants with resistance. This indicates a need for a combination of sequencing methods followed by functional validation to detect metabolic resistance markers. Genome wide association of pooled individuals (GWAS-PoolSeq) has successfully detected candidate genomic regions of specific phenotypes, including variation in pigmentation in *Drosophila* [13]. In *An. funestus*, we recently discovered a duplication of the X chromosome cytochrome P450 *CYP9K1* associated with increased gene expression using this method [9]. Deep sequencing of target-enriched data has successfully been implemented to elucidate mechanisms of insecticide resistance in the dengue mosquito vector, *Aedes aegypti* [14]. Therefore, a GWAS-PoolSeq approach in tandem with targeted enrichment of candidate genomics regions could offer further opportunities to elucidate the complexities of metabolic resistance in *An. funestus*, while also helping to detect causative resistance alleles.

Here, we used a multi-omics approach with a GWAS-PoolSeq and target enrichment with deep sequencing to elucidate the molecular basis of pyrethroid resistance in the major malaria vector *An. funestus.* We show, using *Escherichia coli* heterologous expression, that a highly selected allele of the cytochrome P450 gene *CYP9K1* is driving pyrethroid resistance in East Africa with complete fixation in Uganda. *In vitro* heterologous expression of *CYP9K1* in *E. coli* revealed this P450 capable of efficiently metabolising the type II pyrethroids deltamethrin.

## Results

### 1 Genome-wide association study with pooled mosquitoes to identify allelic variants putatively associated with permethrin resistance

To detect genetic markers associated with permethrin resistance across the *An. funestus* genome, we carried out a genome-wide association study using pooled mosquitoes with binary ‘resistant’ or ‘susceptible’ phenotypes. Insecticide exposure bioassays were performed on susceptible and resistant populations of mosquitoes from two locations (Malawi and Cameroon) representing Southern and Central Africa, respectively. The ‘Susceptible’ mosquitoes were those that died after a short exposure to the insecticide while ‘highly resistant’ mosquitoes survived a long exposure. Genomic DNA from each mosquito was purified and equal quantities from 40 mosquitoes from each set pooled and whole-genome shotgun sequenced. The sequence data obtained for each F_1_ pool were processed for quality control (trimming, pair-end) (Table S1) and aligned to the *An. funestus* F3 FUMOZ reference genome [15]. Allele frequencies were estimated at all variant sites and compared using pairwise and global F_st_ between susceptible/resistant population and a Cochran-Mantel-Haenszel (CMH) test of association.

To identify variant sites with allele frequencies significantly associated with the phenotypes in Malawi [dead (D) after 60 min’ permethrin exposure (n=2 pools) and alive (A) after 180 minutes’ permethrin exposure (n=3 pools)], Cochran-Mantel-Haenszel tests of association and a gene-wise divergence (*F_ST_*) estimations were applied to all bi-allelic variant sites. These estimates were plotted as -log10 P-values Manhattan plots for 1000 SNP sliding-window global F_st_s estimated in the R package poolfstat [16] and Cochran Mantel Tests of association (Figure 1a-b) using Popoolation2 [17]. GWAS Results were consistent between Cochran-Mantel-Haenszel (CMH) tests of association and global F_st_. For both analyses, a twelve megabase-long region of elevated F_st_/-log_10_ p-value is observed between 21 and 33 Mb on chromosome 3 in Malawi. This extensive region is annotated with 765 genes many of which are of unknown function (242) but does include six cuticular genes and one cytochrome P450 (*CYP301A1*). The average F_st_ in this region is 0.018 versus a background of *F_st_* of 0.0005 for chromosome 3. Although a substantial 32-fold difference in F_st_ averages, the absolute F_st_ of this region was low. Furthermore, on inspection of the pairwise *F_st_* plots (Figure S1), this elevated region was observed in “Alive1” and “Alive3” versus dead replicates but not for “Alive2” replicates. Two peaks on Chromosome 2 around positions 95.6 and 97.7 are prominent in the F_st_ results and can also be discerned in the CMH plots (Figure 1). The first region of elevated F_st_ from positions 95,515,427 to 95,668,792 is composed of 40 genes including *CYP9M1* (AFUN015938) and *CYP9M2* (AFUN016005). There were also four cellular retinaldehyde binding proteins, three CRAL-TRIO domain-containing proteins and the remaining 31 genes lacked annotation. The peak around 97.7Mb did not overlap any gene but is downstream of the 3’ end of gene AFUN003294 which encodes an ETS family transcriptional repressor. The only other visually concordant region of potential interest was observed towards the end of the X-chromosome from positions 14.4 to 14.7 mb overlapping four genes including a homolog of *‘single-minded’* (AFUN005600) and an un-annotated gene (AFUN020237) with homology to ‘*stasimon’* where local F_st_/-log10 p-values were highest. Although the SNP with the highest CMH -log_10_ p-value is outside of this region at position 14,172,028.

**Figure 1.**
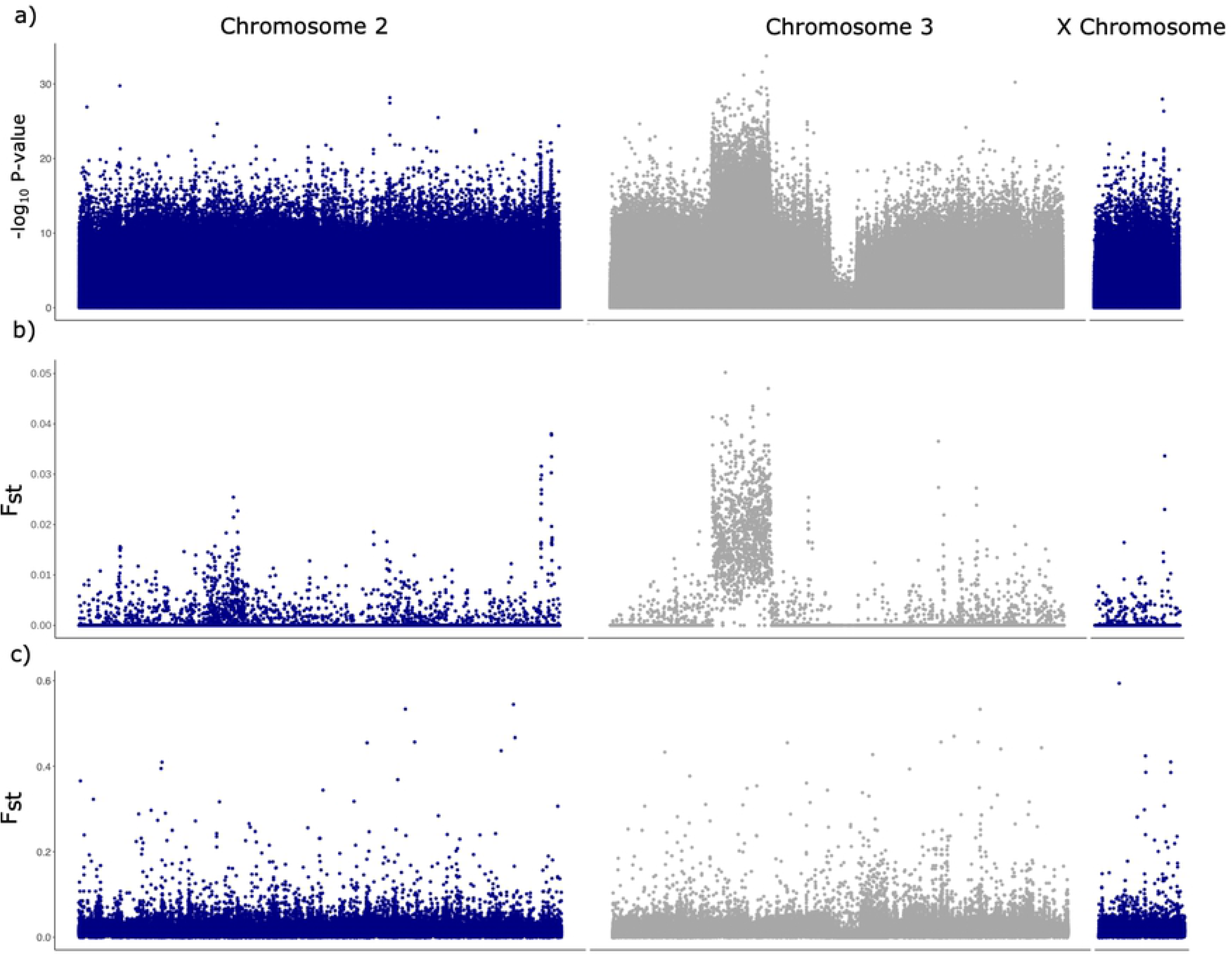
PoolSeq genome-wide analysis between pools of permethrin resistant and susceptible *An. funestus* from Malawi and Cameroon. a) Cochran-Mantel-Haenszel test –log_10_ P-values per SNP calculated in Popoolation 2 in Malawi, b) F_st_ values for 1000 bp windows calculated in poolfstat in Malawi and c) is for Cameroon.

To test if similar results were observed in Cameroon, an F_st_ only analysis was performed as only one Dead and one Alive replicate were sequenced. Background F_st_ values were low and similar for all three chromosomes at 0.015, 0.015 and 0.017 for Chromosomes 2, 3 and X respectively (Figure 1c), although several fold larger background F_st_s in the Malawi data. Outliers were few and did not overlap with those for Malawi. Of those 1000bp blocks with an F_st_ >0.4 only one contained more than 3 SNPs on Chromosome 2 mid-point 67,266,500 with 6 SNPs. This window did not overlap with any genes in the *An. funestus* annotation, and the nearest loci are >5 kb distant in both flanking regions.

Finally, an inter-country comparison was made by poolfstat global F_st_ and Popoolation2 CMH test. In contrast to intra-country comparisons a well-defined peak of differentiation was observed across the *rp1* locus for both analyses (Figure S2). In addition, the X Chromosome was of elevated background F_st_ versus autosomes with average F_st_ of 0.165 versus 0.062 and 0.056 for Chromosomes 2 and 3 respectively. Overall, because of the lack of strong candidate resistance variants detected with this PoolSeq GWAS approach, it was not pursued in other countries, but a fine-scale approach was employed instead.

### 3 Detection of variants associated with pyrethroid resistance using targeted enrichment and deep sequencing

To further detect the polymorphisms associated with pyrethroid resistance, a fine-scale targeted sequencing approach was also used to enrich a portion of the genome using a portion of the genome of individual mosquitoes. The set of genes targeted represent many candidate metabolic resistance loci based on the literature (detoxification genes and previously identified resistance-associated loci). A total of 3,059,528bp of the 1302 sequence capture regions was successfully sequenced in 70 individual mosquitoes (Tables S2, S3 and S4). Mapping and coverage metrics of the targeted sequencing relative to the reference genome were within expectation (Table S3 and S4). The good quality of the target enrichment is also supported by the average base quality of the reads, the alignment score of the mapped reads and the match status of paired ended reads for each sample (Figure S3). Integrative Genomics Viewer (IGV) [18] was used to visually inspect the alignment results showing that in general, sequence capture regions were well covered and lower level coverage was seen between these regions.

A total of 137,137 polymorphic sites were detected across all three countries plus the susceptible FANG laboratory colony. The Malawi samples exhibited lower polymorphisms compared to the reference genome (FUMOZ, originally sampled in southern Mozambique), which is expected as both are from southern Africa. Analysis performed between each country and FANG detected 75,980, 79,095 and 38,380 polymorphic sites respectively in Cameroon, Uganda and Malawi. Detection of the SNPs significantly associated with permethrin resistance was performed firstly using the differential SNP frequency analysis implemented in Strand NGS (Strand Life Sciences, Bangalore, India).

#### Cameroon

Using the frequency-based filtering approach, 92 SNPs out of the 75,980 polymorphic sites were found to be significant between resistant and susceptible field mosquitoes (R-C), 73 between resistant and the FANG (R-S), and 64 between Cameroon susceptible and FANG (Figure 2a). Most of these SNPs were silent substitutions followed by intronic and non-synonymous ones (Table S5; Figure 2b). We considered the best candidate SNPs to be those present commonly between the three comparisons. These common SNPs belong to 16 genes (Figure 2c) including seven cytochrome P450s in the known major pyrethroid resistance QTLs notably *rp1* (*CYP6P4a, CYP6P9b*) on 2R chromosome, *rp2* (*CYP6M1b, CYP6M1c, CYP6S2*) on chromosome 2L, as well as in *rp3* (*CYP9J11*) on chromosome 3L. Further evidence of the association of polymorphisms at *rp1* with the resistance phenotype was the presence of the carboxylesterase gene (AFUN015787) located within this same genomic region. Two cuticle protein genes presented abundant significant SNPs (AFUN009934 and AFUN009937) for all three comparisons. Looking at the nonsynonymous substitutions, two genes showed common amino acid changes for all three comparisons, the P450 *CYP6AK1* (AFUN000518) on the 3L chromosome and the UDP-glucuronosyl transferase (AFUN004976). Analysis of the 92 SNPs significant in the R-C (Figure 2a) comparison revealed that the gene with most non-synonymous substitutions is the immune response gene *APL1C* (four nonsynonymous sites) followed by the carboxylesterase (AFUN015787) (three nonsynonymous sites) (Table S5). There were also other immune response genes such as the chymotrypsin-like elastase and other serine proteases. Such genes were also over-expressed in resistant *An. funestus* mosquitoes in previous studies [5, 11, 19, 20].

**Figure 2.**
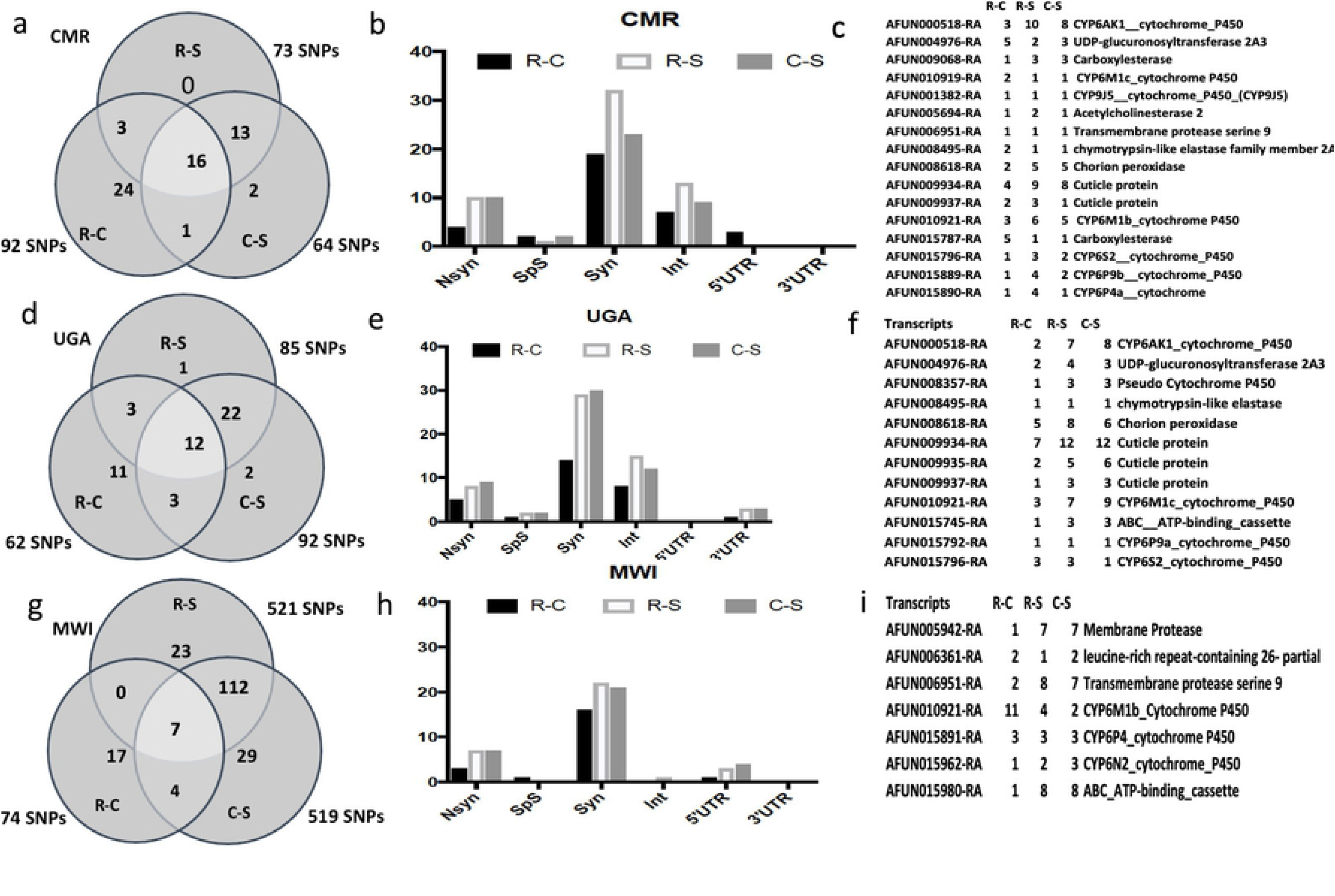
Variants significantly associated with permethrin resistance using SureSelect target enrichment sequencing of specific candidate resistance genomic regions. Using a frequency-based filtering approach implemented in StrandNGS, (a) sets of SNPs significantly associated with resistance were detected in various comparisons between field Permethrin Alive (R) and dead (C) and the lab susceptible strain FANG (S) in Cameroon. (b) Distribution of the significant SNPs between non-Synonymous (Nsyn), Splice Sites (SpS), Synonymous (Syn), intron (Int), 5’untranslated region (5’UTR) and 3’ Untranslated regions (3’UTR) in Cameroon. (b) List of genes with variants significantly associated with permethrin resistance in Cameroon. (d), (e), (f) are equivalent of (a), (b) and (c) for Uganda respectively, as are (g), (h) and (i) for Malawi.

#### Uganda

Using the frequency-based filtering approach, 62 SNPs out of the 79,095 polymorphic sites were found to be significant between Ugandan permethrin resistant and susceptible field mosquitoes (R-C), 85 between resistant and the FANG (R-S) and 92 between Ugandan susceptible and FANG (Figure 2d). Again, as for Cameroon most of these SNPs were silent substitutions followed by intronic and non-synonymous SNPs (Figure 2e). The SNPs present in all three comparisons belong to 12 genes (Figure 2f) including four P450s from the *rp1* QTL (*CYP6P9a* and the pseudo-P450 AFUN008357) and *rp2* (*CYP6M1c* and *CYP6S2*). As for Cameroon, three cuticle protein genes had the most significant SNPs between the three comparisons. Looking at the nonsynonymous substitutions, two genes showed common amino acid changes for all three comparisons, the P450 *CYP6AK1* (AFUN000518) and a cuticle protein (AFUN009934). Analysis the list of the 62 SNPs significant in the R-C revealed that the gene with most non-synonymous substitutions was again the immune response gene APL1C (three nonsynonymous sites) followed by the cytochrome P450 *CY4H19* (AFUN001746) (two nonsynonymous sites) (Table S5). Like Cameroon, there were also other immune response genes such as the chymotrypsin-like elastase and other serine proteases.

#### Malawi

The frequency-based filtering approach detected 74 significant SNPs out of the 38,380 polymorphic sites between Malawian permethrin resistant and susceptible field mosquitoes (R-C) (Table S5), with 521 between resistant and the FANG (R-S) and 519 between Malawian susceptible and FANG (Table S6; Figure 2g). Malawian *An. funestus* are similar genetically to those from Mozambique from which the reference genome FUMOZ was generated, thus, the low diversity seen in this population in comparison to reference genome. Furthermore, due to this similarity to a highly-resistant reference sequence, significant SNPs have been detected assuming a higher frequency in the susceptible than the resistant. Among the common SNPs, the silent substitutions are again predominant followed by non-synonymous and 5’UTR SNPs (Figure 2h). These common SNPs belong to seven genes (Figure 2i) including three P450s from the *rp1* (*CYP6P4a*) and *rp2* (*CYP6M1b* and *CYP6N2*) QTLs. The gene with the most significant SNPs is the P450 *CYP6M1b*. Analysis of the list of the 74 SNPs significant in the R-C revealed that the gene with most non-synonymous substitutions was the P450 *CYP6AK1* (three nonsynonymous sites) followed by the cuticle protein (AFUN009936) (two nonsynonymous sites) and cytochrome P450 *CYP4H19* (AFUN001746) (two nonsynonymous sites) (Table S5). As in Cameroon and Uganda, SNPs in immune response genes were also found in all comparisons.

A second approach consisted in detecting significant SNPs using the t-test in each country provided the following results.

#### Cameroon

Comparing the resistant and the susceptible mosquitoes from Cameroon detected several SNPs with significant allele frequency. However, when considering the Bonferroni multiple testing correction cut-off, these SNPs were not above the threshold when all SNPs were included. However, when the most common SNPs only (present in three or more mosquitoes) were analysed, significant SNPs were detected. The most highly significant SNP was in the cytochrome P450 *CYP304B1* on the 2R chromosome (P=1.7.10^-5^). Analysis of the 29 SNPs with P<0.001 revealed seven SNPs that were also detected with the frequency-based filtering approach above (Table S7; Figure 3a) including a SNP located in chorion peroxidase (AFUN00618), the cytochrome P450 *CYP6M1c* (AFUN010919) on the *rp2* QTL. Some of these 29 SNPs also belong to genes that were significantly over-expressed in resistant mosquitoes such the P450 *CYP315A1* and the glutathione S-transferase *GSTe3*. Three non-synonymous SNPs are detected belonging to the P450 *CYP304B1* (amino acid change: I504V), the chymotrypsin-like protease (AFUN015111) (D476G) and the decarboxylase, AFUN007527 (V169L). A comparison of the resistant mosquitoes of Cameroon to the FANG was performed to detect key regions with highly significant SNPs, which could be associated with resistance, even though population structure could explain most of these differences. Therefore, comparing R-S detected, as expected, very high significance level with top P-value of 7.8 x 10^-48^ corresponding to a cuticular protein gene (AFUN004689). Overall, most of the major genomic regions with the highest significance of SNPs between Cameroon and FANG are found around the pyrethroid resistant QTL *rp1,* and a region made of Zinc finger protein (AFUN015873), as well as a cluster of ABC transporter genes around ABCG4 (Table S8; Figure 3b). This cluster of ABC transporter genes were also detected in the R-C comparison.

**Figure 3.**
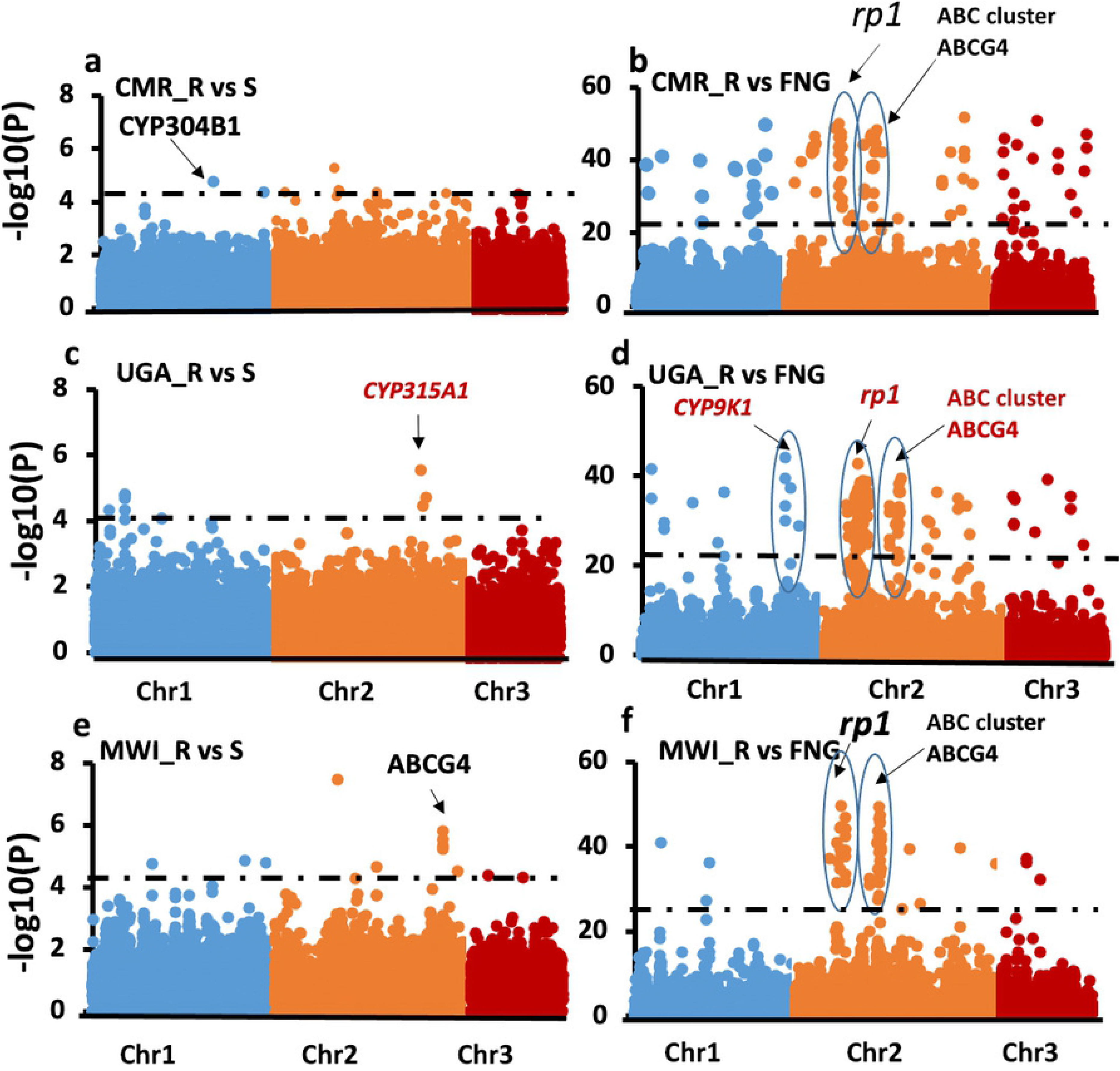
Variants significantly associated with permethrin resistance using an unpaired t-test between the resistant mosquitoes (alive) and susceptible (Dead). (a) Significant variants between permethrin resistant and susceptible mosquitoes in Cameroon (unpaired t-test); whereas (b) is between Cameroon resistant and FANG susceptible strain. (c) is for Uganda Alive and Dead mosquitoes after permethrin exposure while (d) are the significant SNPs between the Uganda Alive and the lab susceptible strain (FANG) and (e and f) are for significant SNPs between Malawi Alive and Dead mosquitoes and versus FANG respectively. SNPs located in the rp1 QTL resistance regions on the 2R chromosomes are consistently associated with pyrethroid resistance. Similarly, a cluster of ABC transporter genes including ABCG4. The black dotted line indicates multiple testing significance level (P=5×10^-5^) for R-C comparisons and (P=5×10^-22^) for comparisons with FANG susceptible strain.

#### Uganda

When the resistant and the susceptible mosquitoes from Uganda were compared (R-C) several SNPs with significant differential allele frequency were detected. As for Cameroon when considering the Bonferroni multiple testing correction cut-off, these SNPs were not above the threshold when all SNPs were included. However, when the most common SNPs only were analysed as for Cameroon some were significant. The most highly significant SNP was in the cytochrome P450 *CYP315A1* (AFUN005715) on the X chromosome (P=2.9.10^-6^). Some of these 53 SNPs also belong to genes significantly over-expressed in resistant mosquitoes such the P450 *CYP315A1* (Table S9; Figure 3c). Six non-synonymous SNPs are detected with some belonging to detoxification genes such as the P450 *CYP6AG1* (K262Q), or to immune response genes such as the transmembrane protease serine 13 (AFUN003078) (H61Y), serine protease 14 (AFUN000319) (N18H), chymotrypsin-like elastase (AFUN015884) (T40K) and the C-type lectin AFUN002085 (L63R). A comparison of the resistant mosquitoes from Uganda to FANG was also performed as in Cameroon detecting as expected a very high significance level with top P value of 2.28.10^-50^ corresponding to an intergenic substitution between the P450 gene *CYP6P9a* and a carboxylesterase gene (AFUN015793) on the *rp1* QTL region (Table S10; Figure 3d). Overall, most of the major genomic regions with the highest significance of SNPs between Uganda and FANG are found around the pyrethroid resistant QTL *rp1* and a cluster of ABC transporter genes around ABCG4 (Figure 3d). Interestingly, a peak of significant SNP corresponds to the *CYP9K1* P450 gene shown to be highly expressed in Uganda and with a marked selective sweep signature around it from whole genome sequencing [9]. Another region corresponded to the argininosuccinate lyase gene which is also highly overexpressed in Uganda compared to FANG.

#### Malawi

Several SNPs with significant differential allele frequency were detected when comparing the resistant and susceptible mosquitoes in Malawi although when considering the Bonferroni multiple testing correction cut-off, some of these SNPs were not above the threshold when all SNPs were included. Among the 59 significant SNPs the top significant was a synonymous substitution in the ABC transporter gene (ABCG4) (A/G, N1347) (AFUN007162) on the X chromosome (P=3.0.10^-8^) (Table S11; Figure 3e). Some of these 59 SNPs also belong to genes significantly over-expressed in resistant mosquitoes such the P450 *CYP6Y1* and the synonymous A/G substitution in the cuticle protein gene AFUN009937 (V48). Four nonsynonymous SNPs were detected with some belonging to detoxification genes such as xanthine dehydrogenase (AFUN002567) (Q799E), or to immune response genes such as the Toll-like receptor (AFUN002942) (V104M). A comparison of the resistant mosquitoes from Malawi to FANG was also performed detecting as expected very high significance level with top P value of 1.7.10^-45^ corresponding to a synonymous substitution in the P450 gene *CYP6P2* in the *rp1* QTL region where a cluster of significant hits is observed (Table S12; Figure 3f). Another cluster of significant hits is also detected around the ABCG4 gene which is also significant between the R-C comparisons (Figure 3f).

### Heterologous expression of *An. funestus* CYP9K1 in *Escherichia coli*

#### Expression pattern of recombinant CYP9K1

A standard P450 carbon monoxide (CO) -difference spectrum was obtained when *CYP9K1* was co-expressed with cytochrome p450 reductase (CPR) in *E. coli*, as expected from a good-quality functional enzyme with a predominant expression at 450 nm and low P420 content (Figure 4a). Recombinant CYP9K1 expressed with a P450 concentration of ∼1.2 nM at 48hr, and a P450 content of 0.93 nmol/mg protein. The membranous P450 reductase activity was calculated as 52.04 cytochrome *c* reduced/min/mg protein.

**Figure 4.**
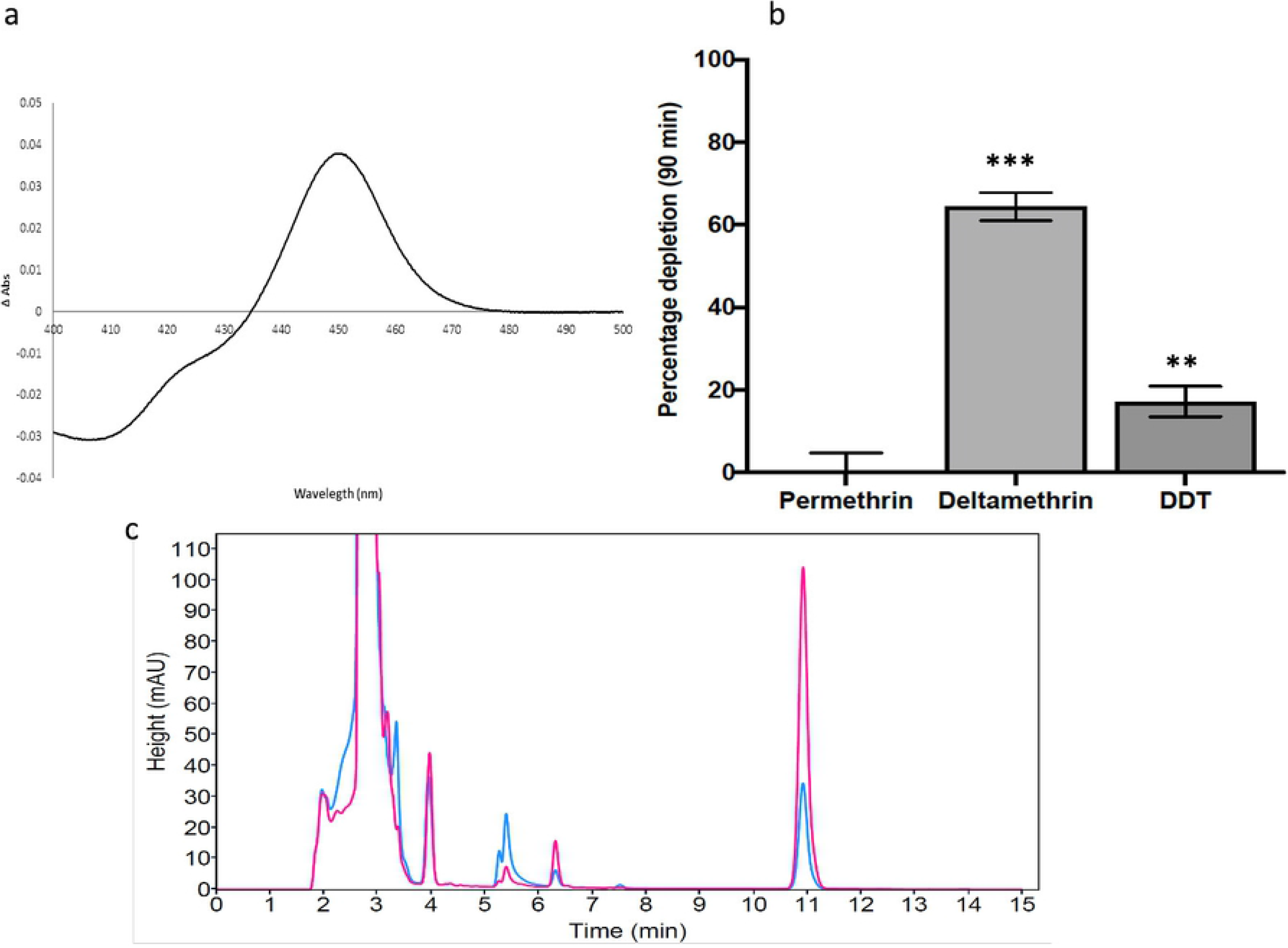
Metabolism of insecticides by recombinant *An. funestus* CYP9K1. a) CO-difference spectrum generated from *E. coli* membranes expressing CYP9K1. b) Percentage depletion of various insecticides (20µM) with recombinant CYP9K1; results are average of 3 replicates compared with negative control (-NADPH); *** Significantly different from -NADPH at p<0.005. c) Overlay of HPLC chromatogram of the CYP9K1 metabolism of deltamethrin, with –NADPH in pink and +NADPH in blue

#### An. funestus CYP9K1 metabolism of insecticides

Recombinant *CYP9K1* exhibited contrasting activity towards permethrin (Type I) and deltamethrin (Type II). While no metabolic activity was observed with permethrin (0.47% depletion), *CYP9K1* depleted 64% (64.37.5% ± 3.44, p<0.01) of deltamethrin in 90min [as determined by the disappearance of substrate (20 µM) after 90 min] compared to controls (with no NADPH) (Figure 4b and c). For DDT, a depletion of only 17% was observed, with no peak for either dicofol (kelthane) or DDE.

### Analysis of *CYP9K1* polymorphism across Africa

#### Comparative analysis of *CYP9K1* polymorphism in resistant and susceptible mosquitoes

A 2,707 bp genomic fragment spanning the full *CYP9K1* gene (5’UTR, 3’UTR, two exons and one intron) was analysed between ten permethrin-resistant and ten susceptible mosquitoes from each of the three countries and from the FANG. Analysis of these 70 mosquitoes revealed 137 substitutions and 72 haplotypes of the 2.7kb gene-body of *CYP9K1* across the continent. When mosquitoes were analysed by country, however, a stark contrast was observed between Uganda and other samples. This was evident for most parameters assessed, notably the lower number of substitution sites in Uganda (35 overall) versus Cameroon (123) and Malawi (42). A similar paucity of haplotypes was observed, with just five haplotypes in Uganda versus 38 and 29 in Cameroon and Malawi, respectively. Not surprisingly therefore, haplotype diversity in Uganda was also very low (0.19) in contrast to Cameroon (0.99) and Malawi (0.97) (Table S13). Similar patterns for Uganda were observed for other parameters including nucleotide diversity (π), this is well illustrated in the plot of haplotype diversity and nucleotide diversity (Figure 5b). Furthermore, Uganda samples exhibited low diversity when compared to the FANG and FUMOZ. Both dead and alive mosquitoes exhibited this low diversity in Uganda (Figure 5a). A similar pattern of reduced polymorphism was seen when considering only the coding region (1614bp) (Table S14) or the non-coding (introns plus UTRs; 1093bp) (Table S15). Analysis of the coding region detected a non-synonymous polymorphism, substituting glycine for alanine at position 454, a mutation which is present in all individuals from Uganda. This G454A change was detected at lower frequencies in Malawi (14/40) and in Cameroon (9/40). An analysis using the Cytochrome P450 Engineering Database (CYPED) [21] reveals that this G454A mutation is between the meander and cysteine pocket, which should impact on activity/catalysis, as amino acids in this region stabilizes the heme structural core and supposed to be involved in interaction with P450 reductase.

**Figure 5.**
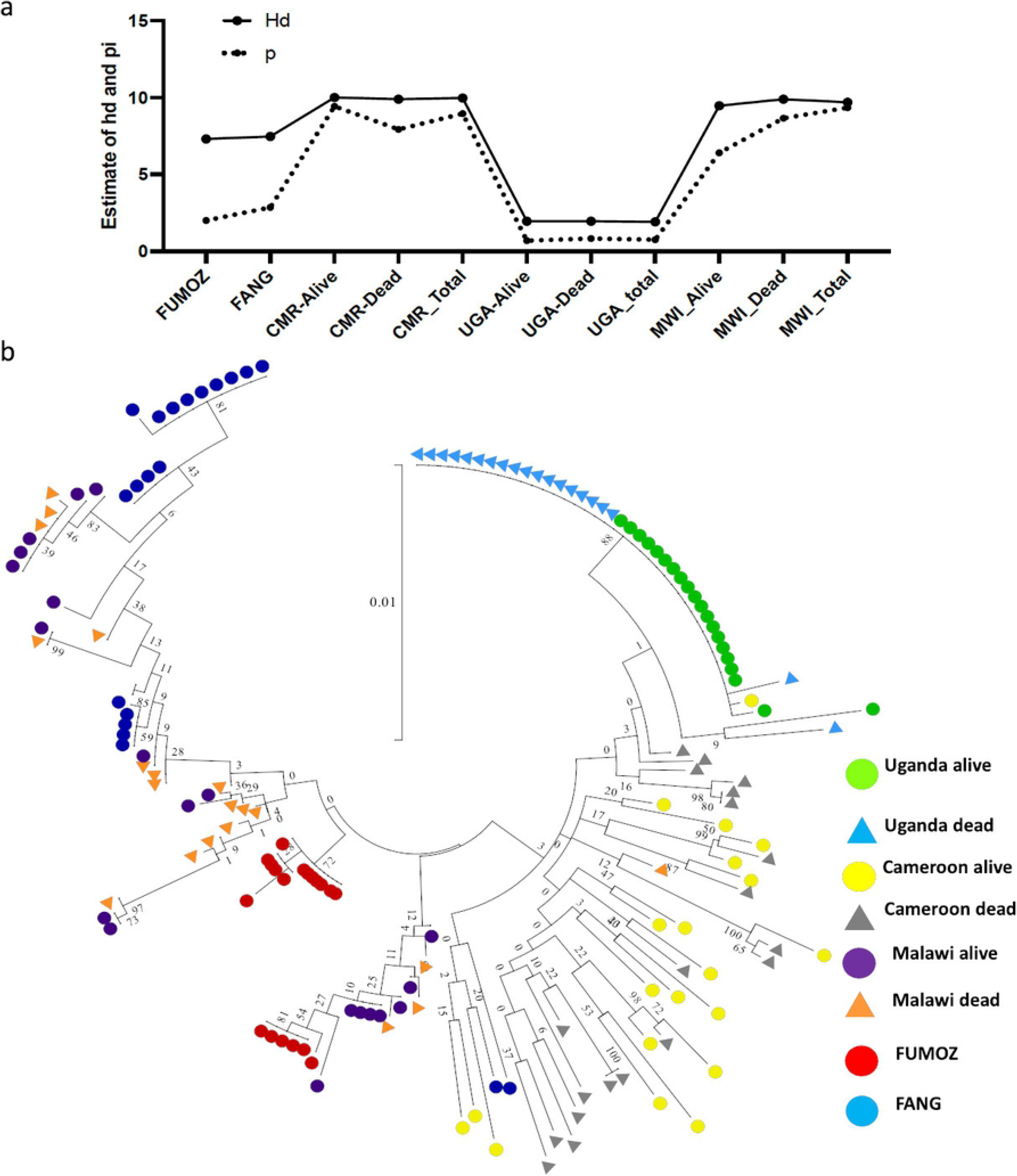
Polymorphisms patterns of *CYP9K1* in Africa. a) Plot of genetic diversity parameters of *CYP9K1* across Africa showing the signature of a strong directional selection of *CYP9K1* in Uganda. H is for haplotype while S is the number of polymorphic sites. b) Phylogenetic tree for *CYP9K1* full-length (2707bp) between Fang and resistant strains of Uganda, Cameroon, Malawi and FUMOZ using SureSelect data.

#### Phylogenetic tree

A maximum likelihood tree of *CYP9K1* sequences supported the high genetic diversity of this gene across the continent with several haplotypes clustering, mostly by their geographical origin (Figure 5b). While mosquitoes from other countries cluster randomly, the majority of those from Uganda belong to a major predominant haplotype (36 out of 40 sequences).

#### *CYP9K1* Haplotype Network

Analysis of the Templeton, Crandall and Sing (TCS) haplotype tree further highlighted the high polymorphism of *CYP9K1* across Africa with many singleton haplotypes separated by many mutational steps (>30 steps) (Figure S4a-b). The predominant haplotype ‘H1’ was nearly fixed in Uganda (32/40) when considering the full-length or also only the coding region (36/40). The fact that this H1 haplotype is shared by both alive and dead mosquitoes suggest that it is close to fixation in this population. In other countries most haplotypes are found as singletons (35 out of 40 in Cameroon; 22 out of 40 in Malawi) supporting the high diversity of *CYP9K1* in those locations in contrast to Uganda. This pattern is similar when only analyzing the coding region (Figure S5a-b) or the non-coding (Figure S6a-b).

## Discussion

As malaria prevention still relies heavily on insecticide-based interventions, it is essential to improve our understanding of the mechanisms driving resistance in malaria vectors to prolong the effectiveness of these tools by implementing suitable resistance management strategies. The present study used a multi-omics approach, and one of these approaches detected that the cytochrome P450 *CYP9K1* is a major driver of pyrethroid resistance in East African populations of the major malaria vector *An. funestus*.

### 1 Genome-Wide Association study with the PoolSeq Approach probably needs more replications

The replicated PoolSeq-based genome-wide association study did not detect significant variants associated with resistance. This is contrary to the usefulness of this method previously in detecting variants associated with natural pigmentation variation in *Drosophila* [13]. Among possible reasons for the lack of sensitivity of this is the poor phenotype segregation in our samples from Malawi. Resistance to insecticide was already relatively high in this population reducing the ability to differentiate between resistant and susceptible. Additionally, increasing the number of replicates could have increased power of detection unfortunately the high resistance level made it difficult to generate sufficient susceptible individuals per location. This was the case for the *Drosophila* pigmentation experiment where more replicates of larger pools of flies were analysed [13], not available to us here as stated above. Despite the low number of candidate hits detected, the elevated region on Chromosome 3 in Malawi does contain six cuticular protein genes (belonging to the RR-2 family associated with the reduced penetration resistance mechanism [22]. This could indicate that the reduce penetration resistance mechanism through cuticle thickening is playing a role in resistance to pyrethroid in Malawi. However, it is noticeable that no hit was detected on the 2R chromosome region spanning the resistance to pyrethroid 1 QTL (*rp1*), which was observed between Malawi and Cameroon. This is likely due to the fixation of selected alleles at these two P450 genes [4], and highlights a drawback in our binary alive versus dead phenotypes as a proxy for resistant and susceptible genotypes. This is similar to the case of knockdown resistance allele L1014F which being fixed in many populations of *An. gambiae* does not correlate with phenotype when using field samples mainly due to the high selection in these samples [23, 24]. The validity of the poolfstat and Popoolation 2 approaches was nevertheless confirmed by the southern (Malawi) versus Central Africa (Cameroon) analysis which detected differentiation at the *rp1* locus. This locus contains the *CYP6P9a* and *CYP6P9b* cytochrome P450 genes, which confer pyrethroid-resistance and are under strong directional selection in southern African populations of *An. funestus* [5, 11, 19]. Although statistically attractive, the replicated PoolSeq offers us little extra over inter-country comparisons of pooled-sequencing as demonstrated by detection of the *rp1* locus here and prior work [5, 9]. Perhaps, a PoolSeq approach using a crossing of resistant strains to susceptible one could provide a more productive platform to detect genetic variants associated with related resistance as implemented in *Aedes aegypti* [25].

### 2 Deep targeted sequencing of genomic regions spanning detoxification genes detects genetic variants of interest

A fine-scale approach combining targeted enrichment and deep sequencing successfully detected variants associated with pyrethroid resistance. This was most evident when comparing resistant mosquitoes to the fully susceptible laboratory FANG strain than when alive and dead mosquitoes from the same location were compared. This low power of detection when comparing samples from the same locality is likely due to high level of resistance inducing a poor segregation between samples. If the high number of significant variants detected between resistant and susceptible strain could be due to a difference in genetic background, the fact that key genomic regions previously associated with resistance were clearly and consistently detected such as *rp1*, revealed the ability of this approach to detect resistance mutations. Indeed, the *rp1* QTL region harbouring a cluster of P450s involved in resistance such as *CYP6P9a/b*, *CYP6P4a/b, CYP6P5* was one of the major loci detected. This could explain why this region was significantly associated with resistance in all regions since at least one gene from this region is over-expressed in each region with *CYP6P5* in Cameroon and Uganda, *CYP6P9a/b* in Malawi [5, 11]. Furthermore, a consistent resistance locus in all three countries when compared to FANG was associated with the ABC transporter gene ABCG4 (AFUN016161-RA) located in the vicinity of two other ABC genes (ABCC4 and ABCC6 as in *An. gambiae*). This highlights the potential important role played by ABC transporters in the resistance to insecticides in general as reported recently [26, 27] and particularly in *An. funestus*. Further work is needed to elucidate the contribution of the gene and variants to the pyrethroid resistance in this species. In Uganda, a significant resistance locus was detected when comparing Uganda resistant to FANG corresponded to *CYP9K1*, in line with country-specific PoolSeq results and RNAseq that show a high over-expression of this gene only in Uganda [5], further support for the likely key role that this P450 gene plays in the pyrethroid resistance in this country [28]. *CYP9K1* has also been implicated in pyrethroid resistant in other mosquito species such as *An. parensis* [29] and *An. coluzzii* [30]. This correlation between RNAseq and targeted sequencing for *CYP9K1* shows that if the phenotypic segregation is wide enough then target enrichment and sequencing could be sufficiently robust to detect variants associated with resistance. Nevertheless, despite narrowing the genomic region associated with resistance to the gene level confirmation of the causative variant requires a further fine scale sequencing of candidate gene and regulatory regions using a classical Sanger sequencing approach followed up by functional genomics such as promoter activity analyses. Without moving to finer scale and functional analyses whole genome studies do not yield the variants needed to design simple molecular diagnostic for resistance tracking of metabolic resistance. An approach we have taken for other metabolic resistance-conferring loci: *GSTe2* [8], *CYP6P9a* [28] and *CYP6P9b* [11].

### 3 *An. funestus* CYP9K1 is a metaboliser of type II pyrethroids

The heterologous expression of *An. funestus* CYP9K1 *(AfCYP9K1)* in *E. coli* followed by metabolism assays revealed that CYP9K1 metabolises the type II pyrethroid, deltamethrin. Recombinant CYP9K1 had a depletion rate similar to those observed for other cytochrome P450s genes in *An. funestus* including CYP6P9b [4], CYP6P9a and CYP6M7 [7], CYP9J11 (CYP9J5) [31] and CYP6AA1 [32] or in other malaria vectors such as CYP6M2 in *An. gambiae* [33] or CYP6P3 [34]. However, the observed *An. funestus* CYP9K1 depletion rate of deltamethrin was twice that for *An. coluzzii* CYP9K1 (64% vs 32%), shown to be conferring pyrethroid resistance in the *An. coluzzii* population of Bioko Island [30] after scale-up of both LLINs and IRS [30]. We hypothesise that the *Af*CYP9K1 allele from Uganda may therefore be significantly more catalytically efficient than the *An. coluzzii* allele selected in Bioko Island. Noticeably, *Af*CYP9K1 did not metabolise the type I pyrethroid permethrin, with no substrate depletion observed after 90min suggesting that *Af*CYP9K1 metabolism is specific to type II pyrethroid. This is similar to previous observations where some P450s could only metabolise one type of pyrethroids. Notably, the CYP6P4 of the malaria vector *An. arabiensis* sampled from Chad was shown not to metabolise type II pyrethroid, deltamethrin, which correlated with susceptibility to this insecticide in this mosquito population [35]. However, we cannot rule out that *Af*CYP9K1 also contributes to type I resistance either through metabolism of secondary metabolites generated by other P450s such as CYP6P9a/b or CYP6P5 also shown to be over-expressed in Uganda [5, 31]. *Af*CYP9K1 could also act through other mechanisms such as sequestration. Considering the very strong selection on this allele established here and previously [9] further studies are needed to establish the extent, if any, of the interaction of CYP9K1 with type I pyrethroids. One possibility is trans-regulation of CYP9K1 as reported for the lepidopteron pest, *Spodoptera exigua* for which trans-acting transcriptional regulators (CncC/Maf) and a cis-regulatory element (Knirps) are both interacting with the 5’ UTR of the P450 gene *CYP321A8*, leading to its upregulation of expression [36].

### 4 A directionally selected *CYP9K1* allele is driving resistance in Uganda

*CYP9K1* is under strong directional selection in Uganda as shown by the polymorphism pattern of this gene in Uganda, with both low numbers of substitutions (35 vs 123 in Cameroon) and haplotypes (5 vs 38 in Cameroon) identified. Strong selection on the *CYP9K1* allele in Uganda is likely driven by the scale up of pyrethroid-based interventions, notably the mass distribution of bed nets. Scale up of bed nets has been strongly associated with the escalation of pyrethroid resistance in southern African *An. funestus* populations [9, 37, 38].

Furthermore, a single haplotype is predominant for *CYP9K1* in Uganda in line with directional selection. Such positive selection is similar to many other cases of cytochrome P450 selected in various insect populations. This is also the case for *CYP6P9a/b* P450s in *An. funestus* for which strongly directionally selected alleles are now fixed in southern African populations [4, 5, 38]. This is also the case for *CYP9K1* in *An. coluzzii* in Mali [39] where an allele has been positively selected in populations post-2006. Similar selective sweeps on P450s have been also reported in *Drosophila melanogaster*, where a single *CYP6G1* allele conferring DDT resistance containing a partial Accord transposable element in the 5’ UTR has spread worldwide [40], [41]. Previous analysis has also shown that the high selection of *CYP9K1* occurs alongside a high level of over-expression related to duplication of the locus of this gene in Uganda [9]. Further supporting selection of an allele with enhanced metabolically efficiency in breaking down pyrethroids. This is supported by the fixation of the amino acid substitution of glycine for alanine at position 454 (G454A). This position is located close to the substrate binding pocket, and we hypothesise that increase the affinity and metabolism of this enzyme for deltamethrin. A similar scenario was seen for *An. funestus CYP6P9a/b* for which both *in vivo* and *in vitro* studies revealed that key amino acid changes (N384S) were able to increase the catalytic efficiency of these enzymes [42]. Further evidence comes from humans for which amino acid changes in *CYP2D6, CYP2C9, CYP2C19* and *CYP2A6* have been shown to affect drug metabolism a low drug metabolism conferred by some alleles while others confer a fast metabolism rate [43]. Similarly, other amino acid changes in the glutathione S-transferase *GSTe2* enzyme in *An. funestus* (L119F) [8] and in *An. gambiae* (I114T) [44] were also shown to drive pyrethroid/DDT resistance in these vectors.

## Conclusion

This study has integrated the combined power of PoolSeq-based GWAS and deep target sequencing of pyrethroid resistant and susceptible mosquitoes with *in vitro* functional validation in *E. coli* of identified candidate genes. We demonstrate that a highly selected *CYP9K1* is driving pyrethroid resistance in Eastern African populations of the major malaria vector *An. funestus*. This result improves our understanding of the molecular basis of metabolic resistance to pyrethroid in malaria vectors and will furthermore facilitate the detection of causative markers to design field applicable diagnostic tool to detect and track this resistance across Africa.

## Materials and Methods

### 1. Design of SureSelect baits

The sequence capture array was designed prior to the release of the *An. funestus* genome assembly, using a mix of *de novo* assembled *An. funestus* transcripts [45, 46] selected from previous pyrethroid resistance microarray experiments [4, 7]. Among these were heat shock proteins (HSPs), Odorant Binding Proteins and immune response genes such as serine peptidases, *Anopheles gambiae* detoxification genes sequences (282 genes) and all target-site resistance genes sequences from *An. funestus*. We also included the entire genomic regions of the major quantitative trait locus (QTLs) associated with pyrethroid resistance which are the 120kb BAC clone of the *rp1* containing the major *CYP6* P450 cluster on the 2R chromosome arm, as well as the 113kb BAC clone sequence for the *rp2* on the 2L chromosome arm. A total of 1,302 target sequences were included (with redundancy). Baits were designed using the SureSelect DNA Advanced Design Wizard in the eArray program of Agilent. The bait size was 120bp for paired-end sequencing using the “centered” option with a bait tiling frequency (indicating the amount of bait overlap) of “x3”.

### 2. Collection, rearing and sequencing of mosquitoes

Two *An. funestus* laboratory colonies (the FANG and FUMOZ) and field mosquitoes from Cameroon, Malawi and Uganda were utilised in this study. The FANG colony is a fully insecticide susceptible colony derived from Angola [47]. The FUMOZ colony is a multi-insecticide resistant colony derived from southern Mozambique [47]. Field populations of mosquitoes representative of Central, East and southern Africa were sampled from Mibellon (6°46′ N, 11°70′ E), Cameroon in February 2015; in March 2014 from Tororo (0°45’ N, 34°5’ E), Uganda [48] and in January 2014 from Chikwawa (16°1’ S, 34°47’ E), southern Malawi [49]. Mosquitoes were kept until fully gravid and forced to lay eggs using the forced-egg laying method [50]. All F_0_ females/parents that laid eggs were morphologically identified as belonging to the *An. funestus* group according to a morphological key [51]. Egg batches were transported to the Liverpool School of Tropical Medicine under a DEFRA license (PATH/125/2012). Eggs were allowed to hatch in cups and mosquitoes reared to adulthood in the insectaries under conditions described previously [50]. Insecticide resistance bioassays on these samples have been previously described [48, 49, 52]. In summary, two-to-five-day old F_1_ females were exposed to permethrin for differing lengths of time to define a set of putatively susceptible (dead after 60 min permethrin exposure for Malawi and Uganda populations, and 20 min for Cameroon) and resistant (alive after 180 min permethrin exposure; 60 min in Cameroon) mosquitoes. The variation of exposure time was associated with the level of resistance in the population.

For the PoolSeq experiment, there were sufficient individuals for two ‘susceptible’ and three ‘resistant’ replicates for 40 individuals each from Malawi and one “susceptible” and one “resistant” replicate from Cameroon. Genomic DNA was extracted per individual using the DNeasy Blood and Tissue kit (Qiagen, Hilden, Germany) and individuals pooled per replicate in equal amounts. Library preparation and whole-genome sequencing by Illumina HiSeq2500 (2×150bp paired-end) was carried out by Centre for Genomic Research (CGR), University of Liverpool, United Kingdom. The SureSelect experiment consisted of ten permethrin susceptible and ten resistant mosquitoes from Malawi (Southern), Cameroon (Central) and Uganda (Eastern) Africa from the set used for the PoolSeq, above. An additional ten mosquitoes from the susceptible FANG strain were also included. The library construction and capture were performed by the CGR using the SureSelect target enrichment custom kit with the 41,082 probes. Libraries were pooled in equimolar amounts and paired-end sequenced (2×150bp) with 20 samples per run of an Illumina MiSeq by CGR, using v4 chemistry.

### 3. Population genomic pipelines

#### 3.1. Analysis of PoolSeq data

The PoolSeq data was analysed in the R package poolfstat [16] and Popoolation2 [17] in order to cross-validate inferences. Both approaches were designed specifically for pooled sequencing datasets. For poolfstat, PoolSeq R1/R2 read pairs were aligned to the VectorBase version 52 *An. funestus* reference sequence using bwa [53]. Output BAM alignment files were co-ordinate sorted and duplicates marked in Picard (http://broadinstitute.github.io/picard). Variant calling was carried out using Varscan (2.4.4)[54], with a minimum variant frequency of 0.01 and p-value of 0.05 and default parameters for other options. Variants were filtered in bcftools (1.9)[55] to remove SNPs within 3 bp of an indel and retain only SNPs for F_st_-based analyses. F_st_ statistics were then calculated from the VCF file with poolfstat. For pairwise intra-Malawi and Cameroon resistant versus susceptible average F_st_ was calculated pairwise between replicates and summarised into non-overlapping 1000 bp windows using ‘windowscanr’ (https://github.com/tavareshugo/WindowScanR/). For all replicates combined analyses of Malawi and Cameroon versus Malawi analyses average F_st_ of non-overlapping sliding windows of 1000 SNPs were calculated within poolfstat.

For Popoolation 2 analyses, a sync file was created from a samtools mpileup (v1.12) and separate comparisons of “Dead versus Alive” and “Cameroon versus Malawi” input to the Cochran-Mantel-Haenszel (CMH) test script “CMH-test.pl”. Only sites with total coverage greater less than 10x and less than the 95^th^ centile for each sample were considered. This test uses multiple independent pairwise comparisons to identify the signals common to all. In this case, the data do not conform to the usual use-case for the CMH test, in which multiple 2×2 contingency tables are stratified by, for instance, location or experiment. Here, independent exposure assays were used to generate the dead and the alive mosquitoes, therefore any pair of samples used to generate a 2×2 contingency table is arbitrary. Using all six possible pairwise combinations of the two Dead and three Alive samples means that the 2×2 tables are not independent of one another and violates the assumptions of the test. This test was run, however, to compare the results to those from tests using independent comparisons. These were six runs of the test made each with two different, independent pairwise combinations of dead and alive samples. Genome-wide F_st_ and -log_10_ p-value plots were created in R using ggplot2 [56] for poolfstat and Popoolation results, respectively.

### 3.2. Analysis of SureSelect data

Initial processing and quality assessment of the sequenced data was performed as for the PoolSeq data and analysed using StrandNGS 3.4 (Strand Life Sciences, Bangalore, India). Alignment and mapping were performed using the “DNA alignment” option against the whole genome (version AfunF1) which was constructed into three chromosomes using synteny from *An. gambiae* [5, 9]. Aligned and mapped reads were used to create a DNA variant analysis experiment. Before variant detection, a SNP pre-processing was performed to reduce false positive calls: (i) split read re-alignment of partially aligned split reads and noisy normally aligned reads; (ii) local realignment to reduce alignment artefacts around indels; and (iii) base quality score recalibration to reduce errors and systematic bias.

All variant types [SNPs, MNPs (multiple nucleotide polymorphisms) and indels] were detected by comparing against the FUMOZ genome using the MAQ independent model implemented in StrandNGS 3.4 and default parameters. A SNP multi sample report was generated for each sample. For each variant, its effect was predicted using the transcript annotation (version AfunF1.4). To identify SNPs significantly associated with permethrin resistance, two approaches were used. Firstly, we used a differential allele frequency-based approach where a variant was significant in relation to permethrin resistance if the supporting read range of the SNP was 35-100% in alive mosquitoes (R) after permethrin exposure and 1-35% in dead mosquitoes (C) (R-C comparison). Both sets of mosquitoes were also compared to the fully susceptible laboratory colony, FANG (S), with significant SNPs having frequency >35% but <35% in FANG (S) in R-S and C-S comparisons. A cut-off of supporting samples range of 5 out 10 was applied to select the SNPs. The second approach assessed the significant association between each variant and permethrin resistance by estimating the unpaired t-test unpaired of each variant between each comparison (R-C, R-S and C-S) and a Manhattan plot of–Log_10_ of P-value created. A SNP frequency cut-off of three or more samples was applied for this approach.

Finally, the polymorphism pattern of the *CYP9K1* gene was analysed across Africa using the SureSelect data. *CYP9K1* polymorphisms were retrieved from the SNP Multi-sample report file generated through Strand NGS 3.4 for each population. Bioedit [57] was used to input various polymorphisms in the VectorBase reference sequence using ambiguous letter to indicate heterozygote positions. Haplotype reconstruction and polymorphism analyses were made using DnaSPv5.10 [58]. MEGA X [59] was used to construct the maximum likelihood phylogenetic tree for *CYP9K1*.

### 4. Heterologous expression of recombinant CYP9K1 and metabolic assays

#### 4.1. Amplification and cloning of full-length cDNA of *An. funestus CYP9K1*

RNA was extracted using the PicoPure RNA isolation Kit (Arcturus, Applied Biosystems, USA) from three pools each of ten permethrin-resistant females from Tororo in Uganda. The RNA was used to synthesize cDNA using SuperScript III (Invitrogen, USA) with oligo-dT20 and RNAse H (New England Biolabs, USA). Full length coding sequences of *CYP9K1* were amplified separately from cDNA of 10 mosquitoes using the Phusion HotStart II Polymerase (Thermo Fisher, UK) (primers sequences: Table S16). The PCR mixes comprised of 5X Phusion HF Buffer (containing 1.5mM MgCl_2_), 85.7µM dNTP mixes, 0.34µM each of forward and reverse primers, 0.015U of Phusion HotStart II DNA Polymerase (Fermentas, Massachusetts, USA) and 10.71µl of dH_2_0, 1µl cDNA to a total volume of 14 µl. Amplification was carried out using the following conditions: one cycle at 98°C for 1min; 35 cycles each of 98°C for 20s (denaturation), 60°C for 30s (annealing), and extension at 72°C for 2min; and one cycle at 72°C for 5min (final elongation). PCR products were cleaned individually with QIAquick® PCR Purification Kit (QIAGEN, Hilden, Germany) and cloned into pJET1.2/blunt cloning vector using the CloneJET PCR Cloning Kit (Fermentas). These were used to transform cloned *E. coli DH5α,* plasmids miniprepped with the QIAprep® Spin Miniprep Kit (QIAGEN) and sequenced on both strands using the pJET1.2F and R primers provided in the cloning kit.

#### 4.2. Cloning and heterologous expression of *An. funestus* CYP9K1 in *E. coli*

The pJET1.2 plasmid bearing the full-length coding sequence of *CYP9K1* was used to prepare the P450 for expression by fusing it to a bacterial *ompA+2* leader sequence allowing translocation to the membrane following previously established protocols [35, 60]. This fusion was achieved in a PCR reaction using the primers given in Table S16. Details of these PCRs are provided in previous publications [7, 35]. The PCR product was cleaned, digested with *Nde*I and *Xba*I restriction enzymes and ligated into the expression vector pCWori+ already linearized with the same restriction enzymes to produce the expression plasmid, pB13::*ompA+2*-*CYP9K1.* This plasmid was co-transformed together with *An. gambiae* cytochrome P450 reductase (in a pACYC-AgCPR) into *E. coli JM109*. Membrane expression and preparation was performed as for [61]. Recombinant *CYP9K1* was expressed at 21°C and 150 rpm, 48 hours after induction with 1mM IPTG and 0.5mM δ-ALA to the final concentrations. Membrane content of the P450 and P450 reductase activity were determined as previously established [62, 63].

#### 4.3. *in vitro* metabolism assays with insecticides

Metabolism assays were conducted with permethrin (a Type I pyrethroid insecticide), deltamethrin (a Type II) and the organochlorine DDT. Assay protocols have been described previously [7, 32]. 0.2M Tris-HCl and NADPH-regeneration components were added to the bottom of chilled 1.5ml tubes. Membranes containing recombinant *CYP9K1* and *Ag*CPR were added to the side of the tube to which cytochrome b_5_ was already added in a ratio 1:4 to the concentration of the *CYP9K1* membrane. These were pre-incubated for 5min at 30°C, with shaking at 1,200 rpm. 20µM of test insecticide was added into the final volume of 0.2ml (∼2.5% v/v methanol), and reaction started by vortexing at 1,200 rpm and 30°C for 90min. Reactions were quenched with 0.1ml ice-cold methanol and incubated for 5min to precipitate protein. Tubes were centrifuged at 16,000 rpm and 4°C for 15min, and 100µl of supernatant and transferred into HPLC vials for analysis. All reactions were carried out in triplicate with experimental samples (+NADPH) and negative controls (- NADPH). Per sample volumes of 100µl were loaded onto isocratic mobile phase (90:10 v/v methanol to water) with a flow rate of 1ml/min, a wavelength of 226nm and peaks separated with a 250mm C18 column (Acclaim ^TM^ 120, Dionex) on an Agilent 1260 Infinity at 23°C. For DDT, a solubilizing agent sodium cholate (1mM) was added as described in [64] and absorption monitored at 232nm. Enzyme activity was calculated as percentage depletion (difference in the amount of insecticide remaining in the +NADPH tubes compared with the –NADPH) and a t-test used to assess significance.

## Availability of data and materials

All genomic datasets are available from the European Nucleotide Archive. Pooled template whole genome sequencing data are available under study accessions PRJEB24379 (Cameroon and Malawi PoolSeq), PRJEB24520 (Cameroon SureSelect), PRJEB47287 (Malawi and Uganda SureSelect [Release date 1^st^ December 2021]) and PRJEB24506 (FANG SureSelect).

## Authors’ contributions

CSW conceived and designed the study, JRM and CSW collected the mosquito field samples. HI, JMR and GDW prepared all samples for genomic sequencing. GDW, CSW and JH analysed pooled-template genomic data. CSW designed the SureSelect baits and analyse the sequencing data. SSI performed the *CYP9K1* metabolism assay and sequence characterisation of *CYP9K1*; LJM, BFT and CDT analysed the *CYP9K1* polymorphism; JH and CSW wrote the manuscript. All authors read and approved the final manuscript.

## Acknowledgements

Pooled-template whole genome sequencing and SureSelect Target enrichment libraries were made and sequenced by the Centre for Genomic Research, University of Liverpool.

## Supplementary Tables

**Table S1.** Descriptive statistics of PoolSeq sequence read data and alignments of permethrin-resistant and susceptible mosquitoes from Malawi and Cameroon.

**Table S2**. Descriptive statistics of SureSelect sequence read data for FANG colony mosquitoes and permethrin-resistant and susceptible mosquitoes from Cameroon, Uganda and Malawi.

**Table S3.** Mapping metrics of the targeted sequencing relative to the reference genome.

**Table S4.** Coverage metrics of the targeted sequencing relative to the reference genome

**Table S5.** Summary of significant SNPs detected between the permethrin resistant (R) and field dead mosquitoes (C) (R-C) in Cameroon, Malawi and Uganda.

**Table S6.** Significant SNPs between Malawi and FANG.

**Tables S7.** List of SNPs significant between the permethrin resistant (R) and field dead mosquitoes (C) (R-C) in Cameroon using

**Table S8.** List of the most significant SNPs between the permethrin resistant (R) mosquitoes in Cameroon and the susceptible lab strain FANG.

**Tables S9.** List of SNPs significant between the permethrin resistant (R) and field dead mosquitoes (C) (R-C) in Uganda.

**Table S10.** List of the most significant SNPs between the permethrin resistant (R) mosquitoes in Uganda and the susceptible lab strain FANG.

**Tables S11.** List of SNPs significant between the permethrin resistant (R) and field dead mosquitoes (C) (R-C) in Malawi.

**Table S12.** List of the most significant SNPs between the permethrin resistant (R) mosquitoes in Malawi and the susceptible lab strain FANG.

**Table S13**. Genetic diversity parameters of the Coding sequences of the *CYP9K1* full sequence (2,707bp) using SureSelect enrichment sequencing.

**Table S14**. Genetic diversity parameters of the Coding sequences only of *CYP9K1* (1,614bp) using SureSelect enrichment sequencing.

**Table S15**. Genetic diversity parameters of the noncoding sequences only of *CYP9K1* (introns and UTR; 1,093bp) using SureSelect enrichment sequencing.

**Table S16**. Primers used for the cloning of CYP9K1 for the heterologous expression in *E. coli*.

## Supplementary Figures

**Figure S1. Pairwise Fst between all Malawi replicates from 1000 bp non-overlapping sliding windows.** a-f) All combinations of resistant (Alive) versus susceptible (Dead) mosquitoes and g-j) resistant versus resistant and susceptible versus susceptible comparisons.

**Figure S2**. **Crude PoolSeq GWAS all Malawi and Cameroon replicates.** a) Cochran- Mantel-Haenszel test –log10 P-values per SNP calculated in Popoolation 2, b) Fst values for 1000 bp windows calculated in poolfstat. SNPs overlapping the *rp1* resistance locus containing the *CYP6P9a/b* cytochrome P450 genes are circled in red.

**Figure S3. Quality control of targeted sequencing:** (A) Average base quality of the reads for one mosquito from Malawi showing the distribution of the base quality score across bases of all reads. (B) alignment score of the mapped reads showing the distribution of reads based on their alignment score. (C) Pie-chart displaying the match status of paired ended reads. This represents the proportion of reads with different read statuses for paired data. (D) The IGV screen showing an overview of the coverage of the some of the targeted genomic regions after SureSelect target enrichment and sequencing.

**Figure S4**. Haplotype network for *CYP9K1* full sequence (2,707bp) using sure select data. a) Alive and dead per population and, b) pooled alive and dead per population.

**Figure S5**. Haplotype networks for *CYP9K1* coding sequence (1,614bp) using sure select data. a) Alive and dead per population and, b) pooled alive and dead per population.

**Figure S6**. Haplotype networks for *CYP9K1* noncoding sequence (1,093bp) using SureSelect data. a) Alive and dead per population. b) Pooled alive and dead per population.

